# GutMeta: online microbiome analysis and interactive visualization with build-in curated human gut microbiome database

**DOI:** 10.1101/2022.09.26.509484

**Authors:** Yiqi Jiang, Yanfei Wang, Lijia Che, Qian Zhou, Shuaicheng Li

## Abstract

**Background:** The human gut microbiome is associated with numerous human diseases. The whole-genome shotgun metagenomics sequencing helps accumulate a massive amount of gut microbiome data. However, few curated integrated platforms are available to explore the vast dataset. Advances in data generation pose new challenges to researchers attempting to analyze, visualize, and reuse published data.

**Result:** GutMeta (human GUT whole-genome shotgun METAgenomics data analysis platform) is a one-stop online human gut metagenomic research platform that integrates a curated database, analyses, and visualizations.

First, we built the Human Gut Metagenomics Database (HGMD), which contained taxonomy profiling and metadata of the metagenomics. HGMD collected the published human gut microbiome samples with whole metagenome shotgun (WMGS) sequencing data and consistently performed taxonomy classification using MetaPhlan3 for each sample. The various related metadata information was curated, and phenotypes were according to the MeSH ID. At this moment, HGMD contains 20,898 samples from 91 projects related to 65 diseases. Embedded tools could help users to explore the samples by keywords. Second, GutMeta provides researchers with user-friendly metagenomics analysis modules, including community diversity calculation, differential testing, dimension reduction, disease classifier construction, *etc*. Then, GutMeta provides corresponding interactive visualizations which can download as Scalable Vector Graphics (SVG), providing high-quality images. Further, GutMeta supplies two additional visualizations for the multi-level taxonomy overview for advanced investigations. GutMeta also supports online editing, including attribute adjustment, recoloring, reordering, and drag-and-drop. Third, GutMeta supports users in building their metagenomics analysis workspaces, including standard profiles uploading and built-in HGMD data import for online customized analyses and visualization.

**Conclusion:** GutMeta offers a solution to improve reproducibility in metagenomic research, with the standardized procedure from input data to downstream analysis and visualization. GutMeta is a free access analysis platform that integrates human gut WMGS sequencing data, nine online bioinformatics analysis and data visualization modules/pipelines, and a customized workspace. GutMeta is avaiable at https://GutMeta.deepomics.org.

## Background

The human gut microbiome is related to multiple human diseases. Initially, researchers focus on the gastrointestinal tract diseases such as inflammatory bowel disease (IBD), irritable bowel syndrome (IBS), colorectal cancer (CRC) [1–7], to metabolic diseases like obesity and type 2 diabetes (T2D) [8–10]. By now, many studies reported immune diseases, cardiovascular diseases, and neurological diseases related to the human gut microbiome. [11–15] In the past two decades, the cost of high-throughput sequencing dropped with metagenomics samples has increased exponentially [16]. Researchers have chosen whole metagenomic shotgun (WMGS) sequencing to study gut microbiome, leading to the enormous human gut microbiome WMGS sequencing data. It brings new insights and challenges in data reusability and results reproducibility. The lack of a standard pipeline causes low effectiveness. Each time researchers need to re-analyze the previously reported samples from raw sequencing data to keep the consistency in processing to avoid system error [6, 7]. At the same time, independent research could not always conclude the same result with different analysis methods, which decreases the repeatability of results and credibility of conclusion. [17]

To deal with the low reusability, some research conducted standard processed human WMGS sequencing samples to standard metadata and taxonomic abundance databases. HumanMetagenomeDB (HMgDB) [18] combines metadata from multiple studies and metagenomes of body parts with collated standardized database but lacks the taxonomic abundance data. The R package curatedMetagenomicData (cMD) [19] collected 90 projects of human microbiome WMGS sequencing sample samples from various body sites with metadata and provided taxonomy profiles and gene family profiles using MetaPhlAn3 and HUMAnN3. GMrepo [20] was an online web database with 17,618 WMGS and 41,285 amplicons of the human microbiome with metadata, but the outdated MetaPhlAn2 conducted taxonomy profiles. Though these three databases declared metadata curation, several shared samples had inconsistent records.

After taxonomic abundance generation, the downstream analysis also reveals no unified among studies. Many toolkits or software packages summarize and integrate analysis methods. Biobackery [21] is a suite of software, including MetaPhlAn3 and HUMAnN3, but it focuses on the upstream profiling of the raw sequencing data to abundance. The R package Vegan [22] had multiple command-line based microbial community analyses with visualization. It focuses on community features, like diversity and ordination. Microbiome helper [23] collects multi-language scripts to use flexible microbiome analysis tools, which have no user interface. The R package animalcules [24] is a new tool that provides analysis and interactive visualization. While those tools usually focus on a specific aspect of analysis and require the users had prior programming knowledge in R/python, which is not userfriendly for the researchers who focus on microbiology. Table 1 summarize the functions and contents of the databases and analysis tools we mentioned. In a word, there is no existing tool for comprehensive analysis of the human gut WMGS sequencing data could be no-programming background friendly and integrated standard processed published data in the analysis.

**Table 1.**
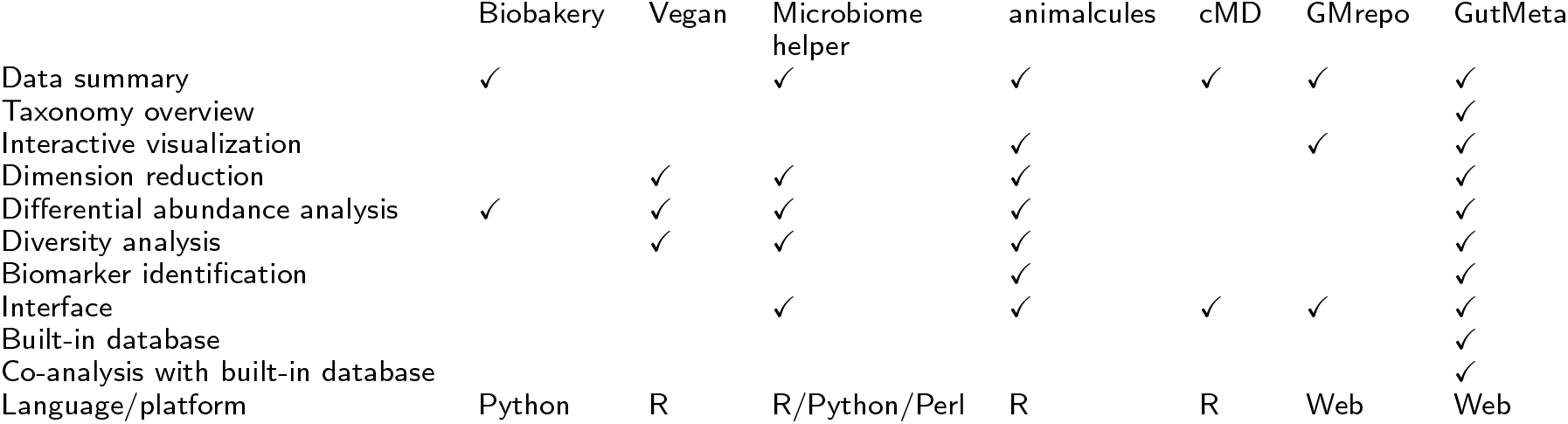

|  | Biobakery | Vegan | Microbiome helper | animalcules | cMD | GMrepo | GutMeta |
| --- | --- | --- | --- | --- | --- | --- | --- |
| Data summary | ✓ |  | ✓ | ✓ | ✓ | ✓ | ✓ |
| Taxonomy overview |  |  |  |  |  |  | ✓ |
| Interactive visualization |  |  |  | ✓ |  | ✓ | ✓ |
| Dimension reduction |  | ✓ | ✓ | ✓ |  |  | ✓ |
| Differential abundance analysis | ✓ | ✓ | ✓ | ✓ |  |  | ✓ |
| Diversity analysis |  | ✓ | ✓ | ✓ |  |  | ✓ |
| Biomarker identification |  |  |  | ✓ |  |  | ✓ |
| Interface |  |  | ✓ | ✓ | ✓ | ✓ | ✓ |
| Built-in database |  |  |  |  |  |  | ✓ |
| Co-analysis with built-in database |  |  |  |  |  |  | ✓ |
| Language/platform | Python | R | R/Python/Perl | R | R | Web | Web |

Here, we present GutMeta, a web-server based one-stop platform for human gut microbiome WMGS sequencing sequencing data. GutMeta consists of the human gut microbiome database (HGMD), online analysis modules, and visualizations. We screened 91 publications that contained public human gut microbiome WMGS sequencing data. We collected 20,898 samples from those publications and curated profiling to abundance. Finally, we integrated the abundance and metadata into HGMD. The abundance and metadata of all samples in HGDB are free access. GutMeta also integrates the metagenomic analysis modules we summarized from previously published works and curated implementation. Users can upload their taxonomy profiling and metadata online for analysis: alpha diversity and beta diversity, dimension reduction, differential testing, and random forest classifier. After researchers obtain the processed data, a well-designed visualization will efficiently help in result evaluation and pattern discovery. Oviz-Bio [25] is a web-based platform that provides interactive real-time online visualization of mutation analysis, widely used in human cancer research. It used Oviz (https://oviz.org), a front-end framework designed for complicated visualization, to draw the diagrams. We developed matching interactive visualizations for analysis based on Oviz. All visualizations provide functions in the interactive information display, online adjustment, and output vector graph in SVG format. At the same time, we designed two visualizations: Overview and taxonomy tree, to let users get a quick overview of microbial community and metadata, and the difference in taxonomic profiling between groups, respectively. With GutMeta, the researcher can analyze their data independently, or with the published samples they are interested in from HGDB without the redundancy and time-consuming steps of downloading and pre-processing the published raw sequencing data. The analysis task would process in the GutMeta cloud server. Users can download and visualize the results online when the analysis is finished. GutMeta is the first online analysis tool with interactive visualization and a built-in database for human gut microbiome WMGS sequencing data. GutMeta provides a possible solution to increase reproducibility and data reusability in human gut microbiome research.

## Implementation

The accessible resource of GutMeta consists of three parts: HGMD database, analysis modules and visualizations (Fig.1). We constructed three parts and integrated them with an in-house web server framework.

**Figure 1.**
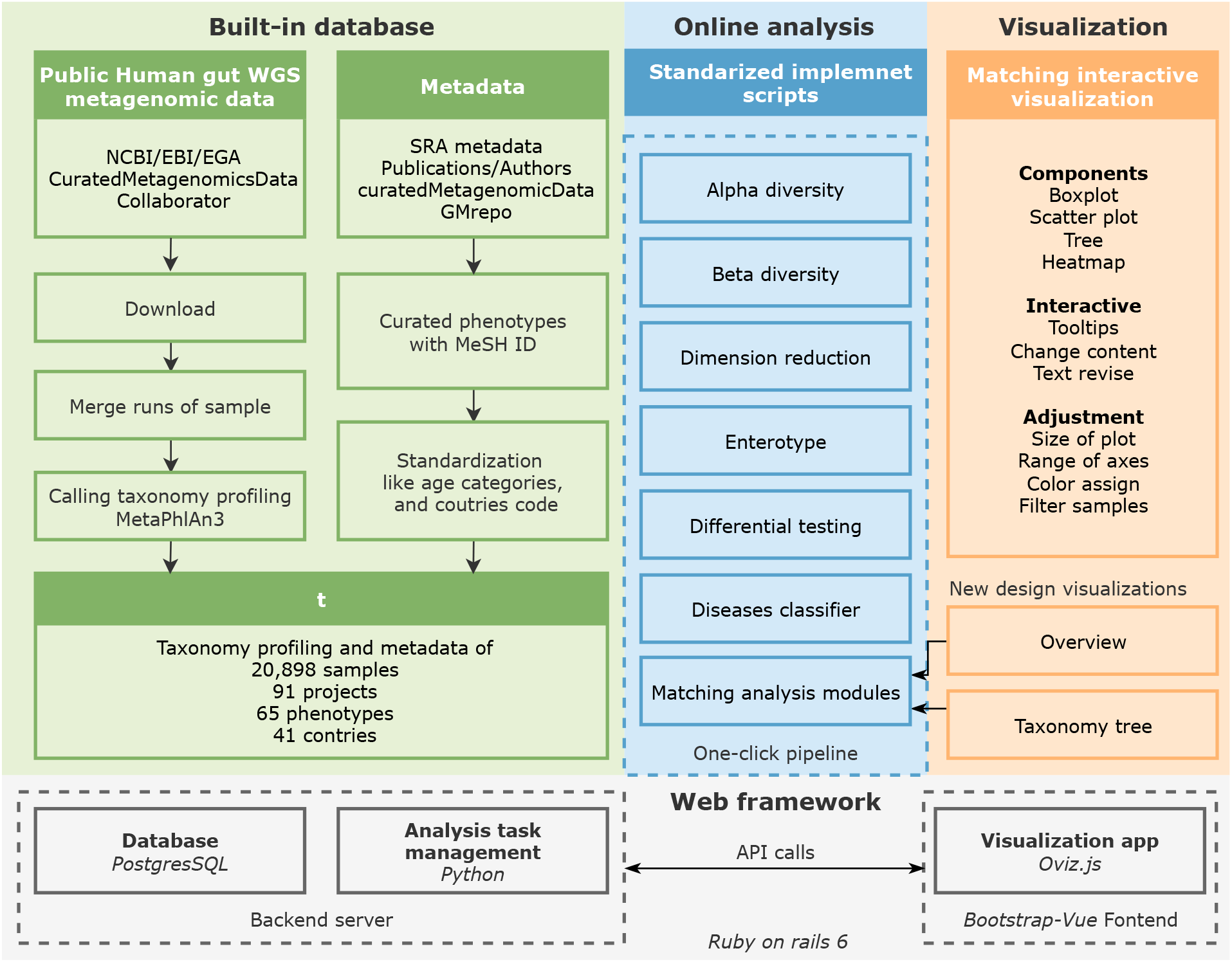
Source content of GutMeta. GutMeta consists of three parts: built-in HGDB with 20,898 samples, standardized analysis modules and pipeline, and interactive online visualizations. Three parts were integrated with a web framework based on Rails.

We summarized the methods and materials we used in the following sections.

### Built-in database

We first collected related publications on the human gut microbiome. The projects we recruited should have published the raw WMGS sequencing data of the human gut microbiome samples on NCBI Sequence Read Archive (SRA) or EBI European Nucleotide Archive. The 91 collected projects are listed on https://gutmeta.deepomics.org/projects.

We downloaded the raw sequencing data in sra format and transformed them into fastq format with SRA toolkits [29]. Compared with the collected stool samples in cMD (version: 3.0.5), we neatened a list of un-collected projects or missing samples. We processed those samples with MetaPhlAn3 [30] using the default parameter for taxonomic assignment. The related metadata was collected from corresponding Bioproject, HMgDB, cMD, and GMrepo. We curated and relabeled the phenotype based on MeSH ID, the country name to ALPHA-3 code based on ISO 3166 international standard, and distinguished samples into different age categories from infant to elder. We left NA to further check if the samples with no metadata provided. After all, we named this database as Human Gut Microbiome Database (HGMD). HGMD currently contains 20,898 samples of 91 projects involving 65 diseases. We organized samples according to the project. Every sample had curated taxonomic abundance with complete taxonomic levels in MPA format and metadata stored in HGMD.

On the database overview page, we provided several statistical results of HGMD. The first one is the distribution of the sample on the world map. After curating merged metadata and manual MeSH labeled, we provided the pie chart of sample distributions for different attributes. We constructed a dataset containing 200 randomly selected samples from healthy adult individuals in HGDB named Healthy _200. Here, we compared Healthy_200 with the sub-datasets containing samples with specific phenotypes stored in HGDB. Only phenotypes that have over 60 samples are included. We applied differential testing module to samples from sub-datasets and Healthy_200, respectively. Finally, we visualized the disease-related taxa with taxonomy tree and bar plots on the Overview page of HGDB. We will introduce the analysis modules and visualizations in the following sections. All statistical resource can be viewed online at https://gutmeta.deepomics.org/database/overview/.

### Analysis modules

We reviewed our collected samples’ corresponding publications and summarized the analysis into six categories: alpha diversity, beta diversity, dimension reduction, enterotype, differential testing and classifier construction. For each analysis, we integrated the commonly applied method and implemented it into curated analysis modules.

Alpha diversity is a vital measurement of the structure of the richness and evenness of microbial communities widely used in ecology and microbiome related research. GutMeta provides several methods to calculate alpha diversity containing Shannon, Simpson, and inverse Simpson using original abundance input. GutMeta also implemented a function to convert abundance to particular reads numbers with data rarefying, letting users can calculate alpha diversity using ACE, chao1, and observed species. With the grouping information provided, GutMeta will calculate the p-value of alpha diversity distribution between groups with chosen method from Wilcox test, t-test, Kruskal Wallis test, and analysis of variance (aov). In the results, GutMeta offers alpha diversity of samples and p-value of comparisons of all taxonomic levels separately from kingdom to species. GutMeta provides a visualization of an interactive box plot of alpha diversity between groups with the p-value.

Beta diversity is an indicator of the dissimilarity or difference between two communities based on distance. GutMeta provides several methods to calculate distance containing Bray-Curtis, Jaccard, euclidean, and Manhatten distances. GutMeta also implemented a function to convert taxonomic annotation in MPA format to the phylogenetic tree. With the function introduced above, GutMeta provides the most popular distance calculation methods used in 16S rRNA analysis, weighted or unweighted unifrac. The beta diversity module provides the same p-value calculation as the alpha diversity module. In the results, GutMeta offers a distance matrix based on the chosen method, and provides an interactive visualization box plot between groups with statistical significant.

The taxonomic abundance of samples usually contains hundreds of taxa. Visualizing all samples in a lower-dimensional space is crucial in gut metagenomics analysis. We provide six methods to reduce dimension on GutMeta: Principal Component Analysis (PCA), Principal Coordinate Analysis (PcoA/MDS), Non-Metric Multidimensional Scaling (NMDS), Independent Component Analysis (ICA), Uniform Manifold Approximation and Projection (UMAP), and t-distributed stochastic neighbor embedding (tSNE). For MDS and NMDS, we provide four distance calculation methods mentioned in the beta diversity module. As a result, the dimension reduction module coordinates samples in all taxonomy levels. GutMeta provides an interactive scatter plot of dimension reduction module visualization. With the metadata of samples, users could visualize the distribution of samples with metadata annotation in scatter point color and shapes and grouped box plots in the chosen two dimensions.

Enterotype is a population stratification approach based on the gut microbiota composition, firstly raised by Arumugam *et al*. in 2011 [8]. The analysis contains two steps: firstly, find the optimal number of clusters by CH index. Secondly, use the Partitioning around medoids (PAM) clustering algorithm to cluster the abundance with a determined cluster number. GutMeta provides PCA and PCoA to visualize enterotypes in low-dimension. The enterotype module on GutMeta integrated the scripts by European Molecular Biology Laboratory [31]. The implementation was based on PAM and clusterSim package in R. The enterotype module provides the CH index, sample clusters, and sample coordinates at every taxonomy level as a result. GutMeta also provides a visualization of the scatter points plot of the enterotype with a highlight ellipse.

The recruited projects in the HGMD majorly focus on the relationship between human diseases and the human gut microbiome. The main clinical application is microbial markers or biomarkers detected from the human gut microbiome. Here, we summarized the biomarker candidates detection analysis in two parts: differential testing and classifier construction. Wilcox rank sum testing is the most suitable statistic method used in metagenomics profiling testing, like the distribution of abundance did not follow any known distribution. We implemented in-house scripts with Wilcox testing in R to find the taxa whose abundance reveals the difference between healthy controls and patients. GutMeta also provides other methods to detect different abundant taxa, like the t-test, Kruskal Wallis test, and aov. The results p-values will be adjusted by the Benjamini-Hochberg method as default or other multiple tests adjustment methods like Hochberg, Holm, Bonferroni, and Hommel.

The differential testing module allows users to set thresholds to filter low abundance and low occurrence taxa to avoid false positive results. With chosen p-value cutoff, the module outputs the significantly different taxa from kingdom to species. GutMeta provides a visualization of this module with compared box plots, automatically descending average abundance sorted taxonomy in all levels. With star symbols to annotated taxonomy fits the strict p-values cutoff set by users, which could be revised on the sidebar editor of visualizations.

After obtained the disease-specific taxa, clinicians excepted to optimize current disease diagnosis. Random forest is the widely-used method to build the disease classifier in the recent gut microbiome related disease research. Here, GutMeta provides an analysis module that can construct and evaluate a disease classifier from the unfiltered taxonomy profiling results based on random forest. It contains all parameters of the differential testing module we mentioned above for users to filter their candidate biomarkers to participate in. It also provides parameters of the fold and replicates number in the cross-validation in feature selection of the construction of classifier. The Classifier module generate the receiver operating characteristic (ROC) curve of the classifier, candidate biomarkers evaluation benchmark and samples prediction result. GutMeta also provides a custom complex visualization for the classifier module. We will describe this in the next section.

### Real-time interactive visualizations with powerful editors

We collected the most frequently used plots for metagenomic analyses based on our developed analysis modules. For each analysis module, GutMeta provides a corresponding interactive visualizations. Every visualization have interactive functions like tooltips (e.g show the element targeted sample), change content (e.g change the taxonomy level of visualization), revise text (e.g revise the graph title, x-axis label), change the size of plot, change the range of axes, customize colors and drag-drop legends at everywhere is applicable. Besides the box plots, scatter points plots and heatmap plots, we designed three complex visualizations with multiple inputs.

#### Overview

Researchers used to draw multiple separate graphs to investigate the abundance distribution, differential testing, and metadata of samples, which would discard the correlation between taxonomy and metadata. Therefore, we designed the ‘Overview’ visualization inspired by the ‘Landscape’ visualization used in cancer research [25]. ‘Overview’ contains taxonomic composition per sample, overall abundance distribution, differential taxa comparison, and metadata of samples. With these crucial plots in one diagram, the ‘Overview’ graph interprets the data more directly and comprehensively. The input files contain the taxonomy profiling and metadata of samples. The fixed part includes the heatmap of taxonomic abundance in samples as the basic structure. The abundance in the heatmap is automatically log normalized. The top stacked bar plots show the microbial composition by sample, while the bottom annotation bars consist of clinical phenotypes, anthropometry information, and artificial grouping of samples. Both categories variable and continuous variable are accepted. The right-hand box plots show the abundance distribution in the two grouped samples.

The sidebar editor provides options to reorder and filter samples, reorder taxonomy, reorder metadata and metadata categories and set colors. In addition, we offered a sorting editor where users can select multiple metadata as criteria to sort the overview diagram, making it easy to explore the correlation between taxonomy and metadata. Users can drag the legends to relocate them and hover on a node, for example, the heatmap grid, to view the information tooltip. Demo visualization is a human gut microbiome of CRC related research in Hong Kong [26] (Fig. 2).

**Figure 2.**
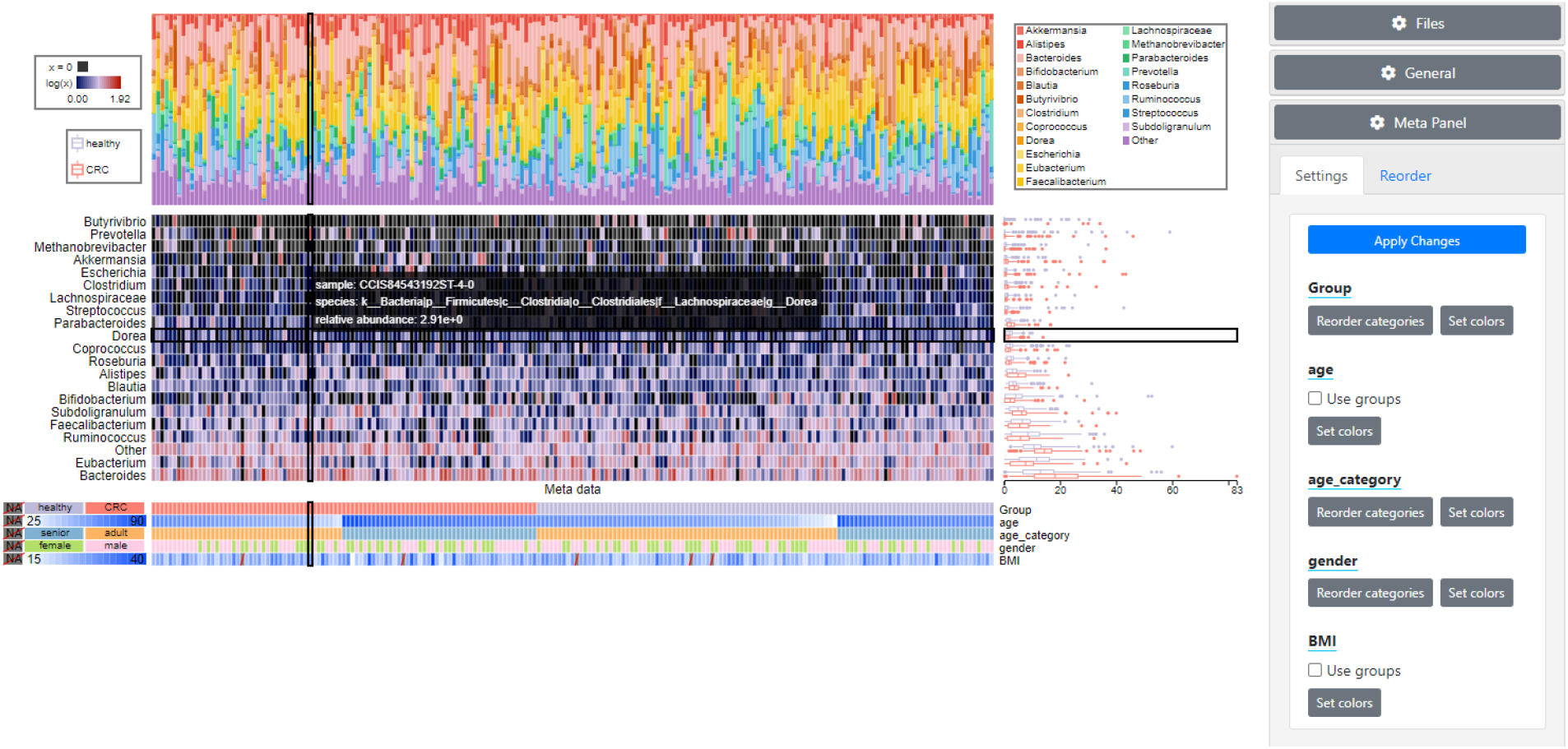
Overview visualizations of Hong Kong CRC cohort [26]. A visualization contains the main display region and an editor on the right hand. Overview visualization contains a heatmap of abundance, a stacked bar plot of microbial composition on the top, compared box plots of the groups, and a metadata heatmap at the bottom. The tooltip shows the pointed sample’s detailed information and highlights the related parts in four panels. The editor can interactively adjust colors and reorder samples and metadata categories.

#### Taxonomy tree

Differential testing of taxonomy abundance between groups is widely-used in metagenomics case-control pattern research. However, compare the abundance in every taxonomy level respectively, ignoring the phylogenetic relationship among levels. Kishikawa *et al*. raised a way to visualize the difference based on a phylogenetic tree [32]. GutMeta provides a ‘Taxonomy tree’ visualization as an interactive optimized version of their design. The ‘Taxonomy tree’ visualization shows profile distribution along the phylogenetic tree with highlights significantly taxa between groups.

The ‘Taxonomy tree’ view contains a phylogenetic tree from kingdom to species. The nodes represent the taxa among samples and the size of nodes is determined by the average relative abundance of the corresponding taxon. The color of nodes is based on the taxa annotation at the class level by default. The branch with highlight color represents the taxonomy enrichment in the corresponding group. The outer layer shows the species name with a rectangle around the significantly different ones. Additionally, the visualization highlighted the significantly different taxa for case-control strategy samples and bordered the species’ names. The flat version of the ‘Taxonomy tree’ also provides the comparison box plots of abundance distribution of two groups.

The sidebar editor provides options to switch the taxonomic level for nodes color, lowest branch level to display, highlight distinct taxa names or not, change the q-values threshold and transform to a flat version. Demo visualization based on IBS-related samples from Nielsen *et al*. with radial form based on species level and flat form based on class level [27]. (Fig. 3)

**Figure 3.**
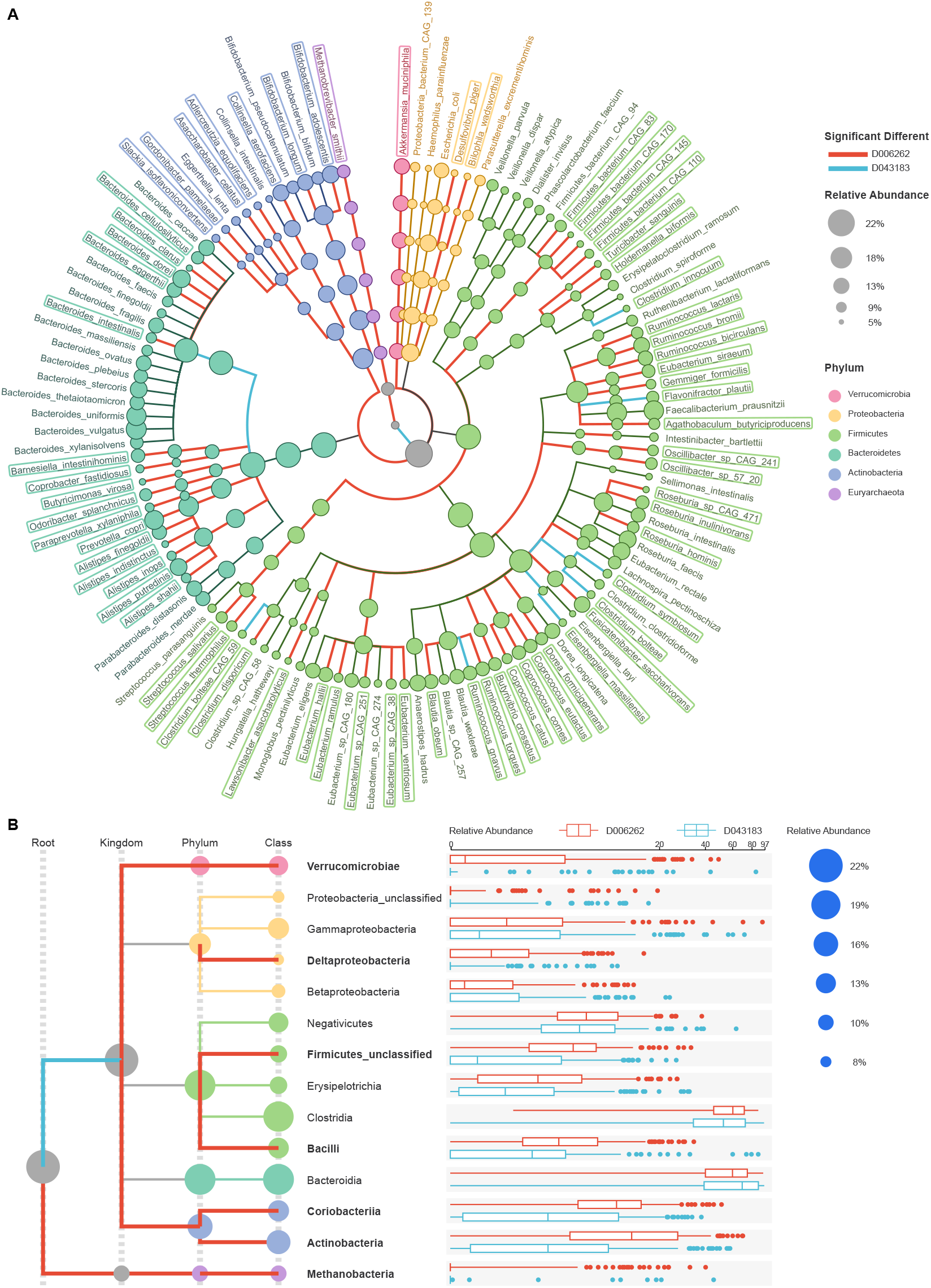
Taxonomy tree of IBS and healthy human gut microbiome comparison [27]. (A) The Taxonomy tree contains taxonomy profiling of two groups of samples with the phylogenetic taxonomy tree from kingdom to species. The background color of tree branches represents different taxonomy. The highlighted branches in red and blue represent the taxonomies significantly different between provided two groups. The outer names are species names, and the significant different species have rectangles around their names. The nodes of the tree reveal the average abundance of the corresponding taxonomy among all samples. The size of the node represents the mean relative abundance of taxonomy. Red lines and blue lines determined the enriched group of significantly different taxonomy. Here groups represent healthy controls and patients. (B) The flat version of taxonomy tree. It contains excess comparison box plots of abundance distribution of two groups.

#### Classifier

The ‘classifier’ visualization displays six graphs the most common plots used in classifier construction and evaluation analysis [28, 33]. The first two graphs are the mean decrease accuracy and the mean decrease Gini of all participated species or genera in crossvalidation and sort in decreasing. The taxon is automatically sorted and the candidates selected as the biomarkers are highlighted. As we can determine the number of biomarkers as described in Qin *et al*. [3] shown in the third graph. The third line chart shows the cross-validation errors with an increasing number of variables selected in independent duplicates. The red line indicates the chosen number of variables. The other three graphs show the classifier’s performance in the training set. We obtain the probability of disease of all samples through the constructed classifier and visualize their distribution in two groups, usually cases and controls, using box plots on the fourth graph and scatter plots increasing on the fifth graph. The last one is the ROC with a 95% CI range of the classifier in the training set with AUC and the 95% CI range noted on the graph.

Here we utilize the flexible of Oviz to make a publication-friendly design. We display the species name on the bar to keep the plots aligned. We color the picked species bars in light pink and the rest in light grey for easy observation. Users can drag the note text to adjust its position in the sensitivity plot and double-click it to edit the content. Users also could customized edit the colors, plot size, and label font size in the editor, and the changes will be applied to all the plots. Demo visualization is myasthenia gravis (MG) related samples from Liu *et al*. [28]. (Fig. 4)

**Figure 4.**
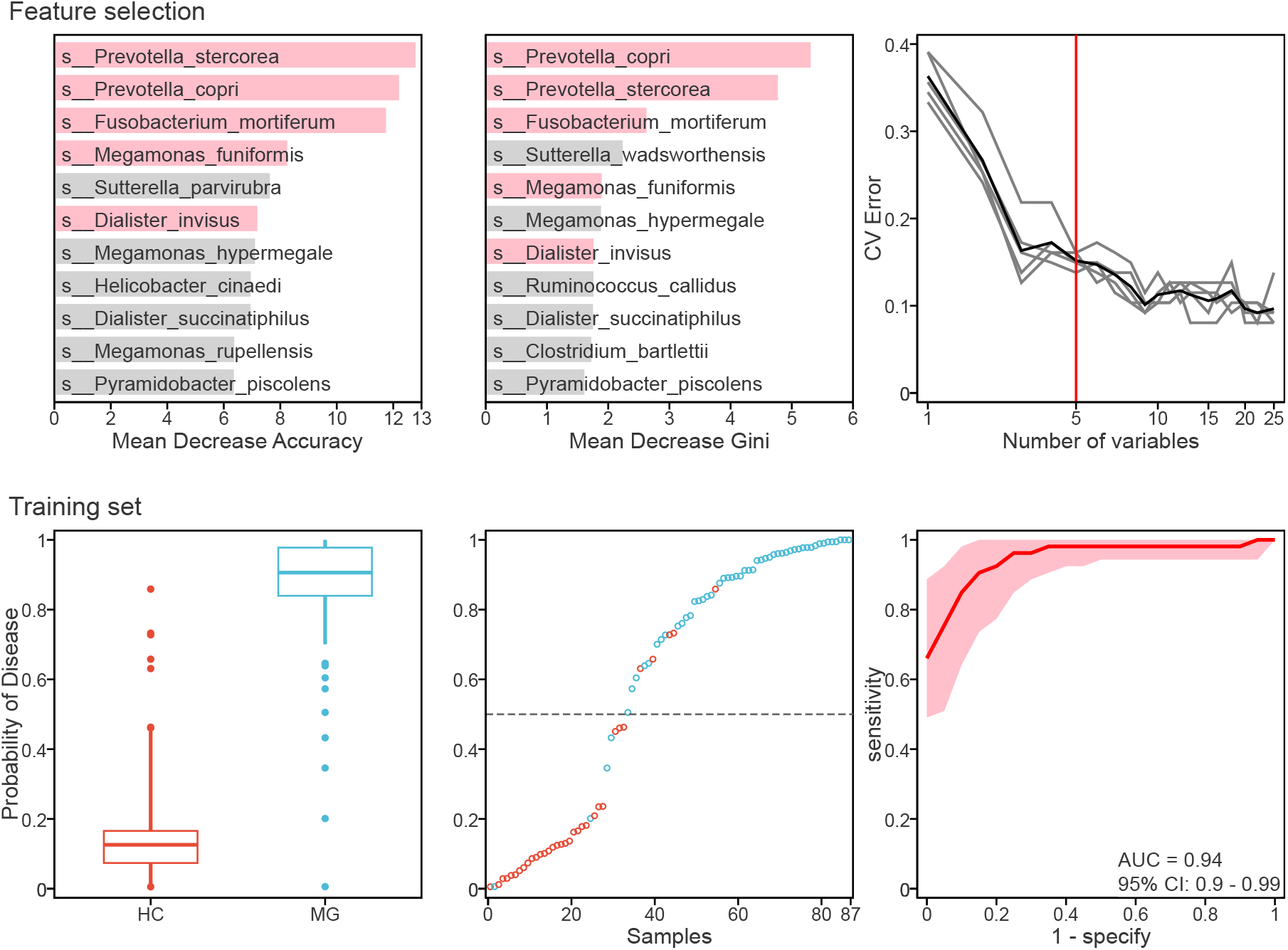
Classifier visualizations of MG patients and healthy controls (HC) [28]. For feature selection, from right to left show mean decrease accuracy of features, mean decrease Gini of features, and number of features had been selected with cross-validation error. Validation in the training set show the disease probability of samples in training set grouped by diseases situations in boxplot and along the probability rank in scatter plot. The last graph is the ROC curve of the classifier, noted with AUC and 95% CI range.

### Inputs standardized and pipeline

We curated the input format of all modules to use abundance in MPA format and metadata. GutMeta also provides the analysis modules to convert standard inputs to the ‘Taxonomy tree’ and the ‘Overview’ visualization’s inputs. All modules contain an input check function to avoid format errors, void records, and unmatching samples between inputs. It will check users’ input and decodes the other format to ASCII format, which is compatible with utf-8. This function makes our analysis function compatible with more input formats and saves users’ time.

Then we construct an all-in-one pipeline containing all the analysis modules mentioned above. Users only need to run with one click and wait to gain all analysis results and visualizations. Up-to-date used analysis scripts can be found at https://github.com/deepomicslab/GutMeta_analysis_scripts.

### Web framework

We constructed GutMeta with Ruby On Rails (version 6.0.2, https://rubyonrails.org), Vue.js (http://vuejs.org), and PostgresSQL (version 9.6, https://www.postgresql.org/). We used Oviz.js, a component-based JavaScript library for data visualization in browsers, to implement all the online data visualizations. In terms of software architecture, GutMeta can be divided into three components: A frontend interface where users can view the HGDB database, upload data for analyses, and view interactive visualized results. A backend server reads and writes the PostgreSQL database server and submits analysis tasks to the task management server through API calls. A standalone task management server written in Python 2.7 that queues and executes the analysis tasks.

## Usage and utility

Users can access GutMeta through the browser on Windows, Unix, and Mac OS. With the navigation bar, users can choose from the following functions: Analysis (run the analysis modules/pipeline, browse demo analysis results), Database (browse samples and projects, download samples metadata and profiles, filter samples to the personal dataset), or visualization (view demo visualization, visualize without analysis). In our design, we divided the users of GutMeta into three groups based on their data: users with gut microbiome dataset, users with interested phenotype keyword, and users with analysis results for visualization. We will introduce the usage and utility of GutMeta with steps shown in Fig.5.

**Figure 5.**
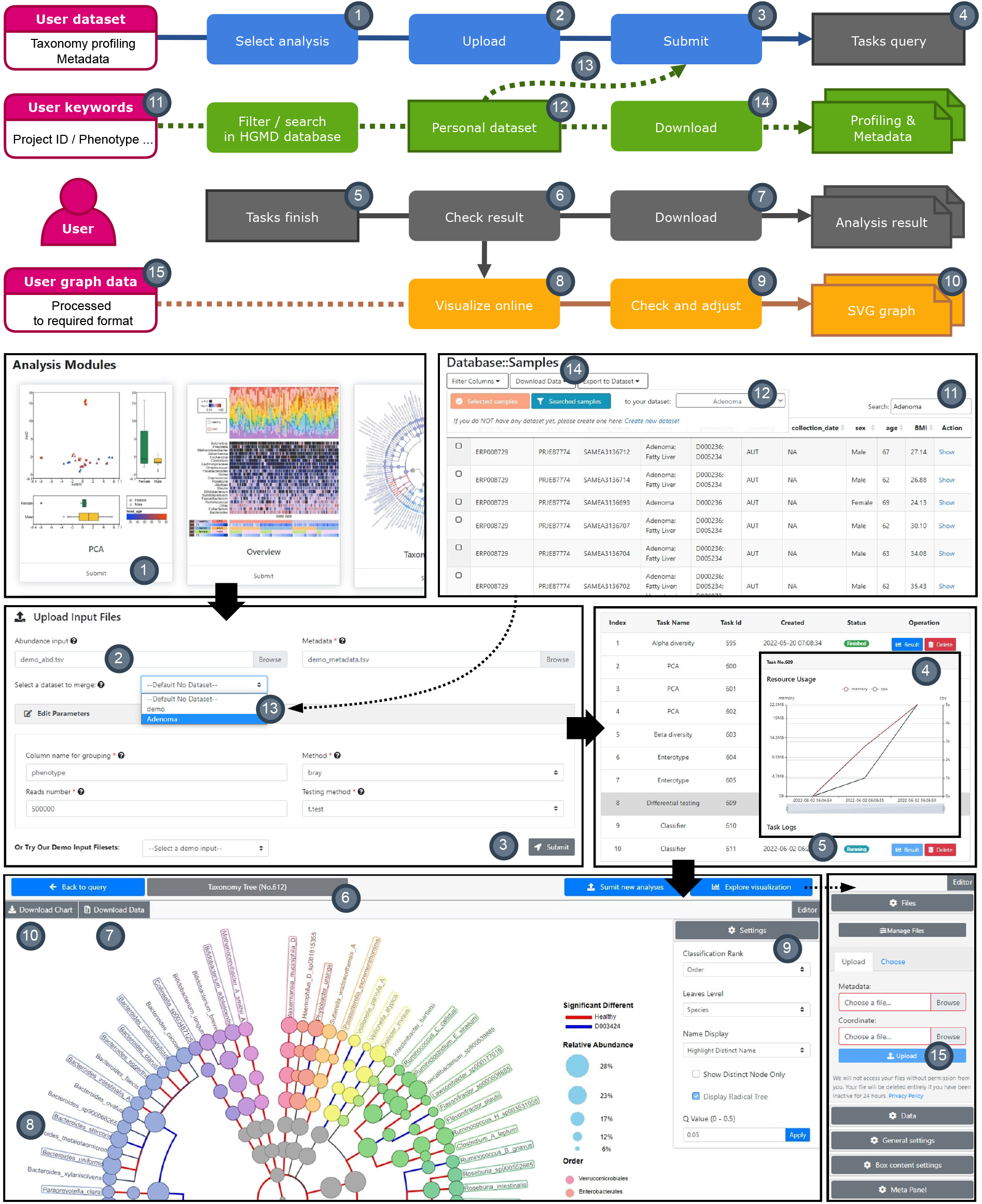
Usage and utility of GutMeta. Number in the grey nodes represents steps in the usage.

Users with their data set could directly analyze them with GutMeta analysis modules. First, users choose the module or pipeline they want to execute on the analysis categories page (step1). Second, upload the abundance and metadata (step2) and submit (step3). Then, users can find their tasks on the job query with real-time computing resource usage (step4). If the task is successfully finished (step5), the user can find the corresponding result of their analysis both in files and visualization. The all-in-one pipeline has the same submit route as modules but will display results module by module (step6).

Afterward, users can view the submitted task on the query page and click the ‘result’ button to open the visualization page. Users can download the result of the module in a compressed tar file (step7), including the data shown in the visualization and the complete evaluations in feature selection after the analysis task is finished correctly. The visualization page contains the main display region (step8) and a sidebar editor (step9) on the right where users can interactively adjust the visualization, detailed adjusted options we have described in the above section. Finally, users can click the Download Chart button to download the vector graph in SVG format (step10).

GutMeta enables users to construct a customized subset of HGDB with their interested keyword. On the sample information page, users could search keywords to filter samples. Here we use “adenoma” as an example (step11). If the user would like to do further analysis online with selected or searched samples, they could export samples to an existing dataset or create a new dataset (step12). On the page of the customized dataset, the user can download the abundance and metadata, and add or delete samples. We also provided three options for users to download the abundance and metadata from the HGMD: selected samples, searched samples, and all samples (step13).

The customized dataset can be used as the input of analysis modules independently or joined analysis with uploaded samples. GutMeta will automatically send abundance data of the data set to the analysis module. GutMeta automatically achieves the combination step. Thus, there is no need for the user to download the dataset and combine it with personal data. Then, the user only needs to select a dataset on the submission page where the user uploads data (step14).

Moreover, if unsatisfied with our analysis results, users can also conduct third-party analysis and upload the files with the validated format on the visualization page (step15). Users can find the file requirements in the User manual section at the bottom of the visualization page and download the demo data in the top tool-bar. Detailed user manual and input file requirements of visualizations can be viewed on each visualization page in GutMeta.

## Results

To illustrate the usage of GutMeta, we include two examples of applying GutMeta in public datasets. The first one contains 100 randomly selected skin microbiome samples from CuratedMetagenomicsData and 100 randomly selected gut microbiome samples from HGDB. The second one is an IBS dataset in HGDB, with 248 samples from healthy individuals and 148 samples from IBS patients. We focus on the microbial community results in the skin-gut dataset and the classifier results in the IBS dataset to reveal how GutMeta works with outer samples and its actual application in human diseases related samples. We set the results of these datasets as demo projects of GutMeta. All results and visualization can view online on the GutMeta demo https://gutmeta.deepomics.org/submit/task-demo with task IDs: 646, 678 and 805.

### Example 1: Overview of the taxonomic composition in microbiome difference between human skin and human gut

The dataset consists of 100 skin microbiome samples from cMD and 100 gut microbiome samples from HGDB. All samples are randomly selectedfrom healthy adult individuals. We applied overview, alpha diversity, beta diversity, and dimension reduction analysis modules to this dataset on the GutMeta.

To check samples’ taxonomic composition directly, we applied the Overview analysis to visualize the abundance at the genus level with samples’ metadata, containing samples group, age, and gender. With the interactive online adjustment of the GutMeta visualization, we sort samples with multiple conditions: by group, descending abundance of *Bacteroides*, and descending abundance of *Cutibacterium*. We spot the huge composition difference between skin and gut microbiome samples. In the top 20 abundant genera, contains well-known gastrointestinal colonizers, like *Bacteroides*, *Eubacterium* and *Prevotella*, and skin niche-specific *Cutibacterium* (Fig.6A).

**Figure 6.**
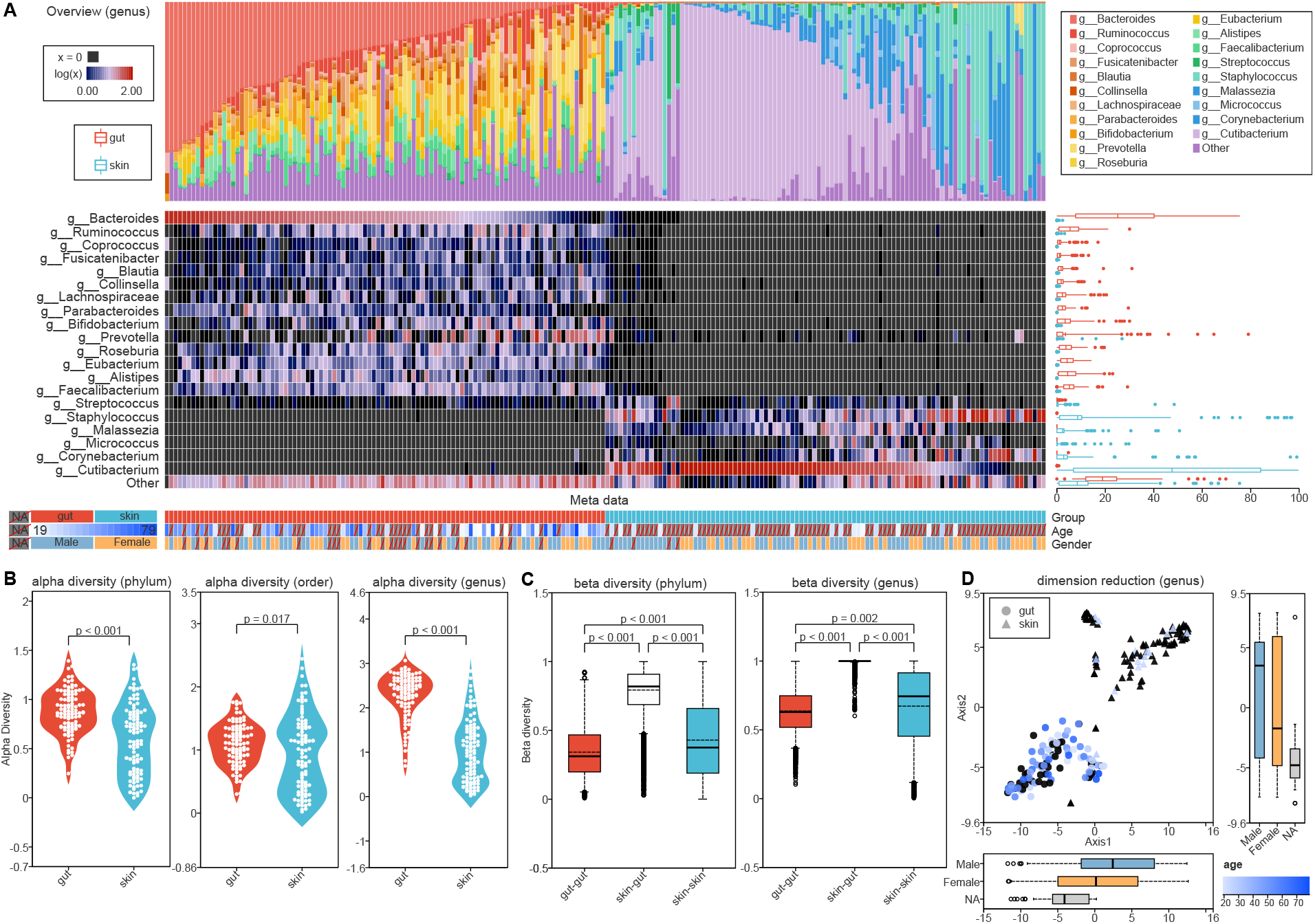
Microbiome composition comparison between gut and skin samples. **(A)** Overview visualization. **(B)** Alpha diversity visualizations in phylum, order and genus. **(C)** Beta diversity visualizations in phylum and genus. **(D)** dimension reduction visualization based on t-SNE.

We applied alpha and beta diversity to this dataset to investigate the community diversity between gut and skin microbiomes. We used the Shannon index to evaluate the alpha diversity among different taxonomy levels. Samples from the gut had significantly higher alpha diversity across all taxonomic levels, while we noticed the difference decrease in order level (t-test, p < 0.05). Compared to the distribution of alpha diversity across samples, we found the cause was the high alpha diversity of five skin samples (Fig.6B). For beta diversity, we used Bray-Curtis distance to evaluate the distance inter-group or intra-groups. The visualizations showed the mean and median distances between groups are higher than two within groups distance. The significantly different beta diversity suggested (t-test, p < 0.001) samples are more similar to the samples in the same group. Skin samples have a higher heterogeneity than gut samples, meaning they have more individual dissimilarity. We also noticed the distance between groups had a decreased variable at low taxonomic levels like genus (Fig.6B).

Ultimately, we applied the dimension reduction method t-SNE to display all samples. We noticed the samples from two habitats clustered in two groups except for two skin samples SAMEA3449372 and SAMN11064157. We could determine those two samples with the interactive tooltips of GutMeta visualization. Back to the overview visualization, we found those two samples had a high abundance in the Other category. At the same time, the box plots reveal the distribution in two dimensions of different gender had no significant difference, which means gender is not related to the microbiome composition difference between gut and skin samples.

### Example 2: CRC dataset re-analysis to construct CRC classifier based on the gut microbiome

In HGMD, we collected 681 samples of CRC patients (MeSH ID: D015179) from 10 independent projects. Previous studies raised young-onset CRC to have distinct clinical and pathologic characteristics [34], we discarded samples from patients younger than 45, and the rest 549 samples labeled old-aged or elder in HGDB participated in the analysis. HGDB contains 3,369 samples from healthy individuals (MeSH ID: D006262) over 45 years old. To balance the sample number, we randomly selected 600 samples from healthy individuals to join the analysis. We applied the analysis modules of the taxonomy tree, differential testing, and classifier to this dataset.

By taxonomy tree, we first investigate the microbiome composition of samples with different disease states. At the phylum level, *Bacteroidetes* significantly enriched in CRC patients. In contrast *Firmicutes* significantly enriched in healthy individuals as they are the top two abundant phyla shown in visualization (Wilcox test, BH adjusted *p* < 0.0001). The taxonomy tree raised a new way to interpret the abundance difference in different taxonomic levels. For example, we noticed the enriched of phylum *Bacteroidetes* in CRC majorly contributed by species *Bacteriodes plebeius, Bacteriodes thetaiotaomicron, Parabacteroides distasonis* and *Parabacteroids merdae*. While, for healthy enriched phylum *Firmicutes*, approximately half of species (26/56) also enriched in healthy individuals, 11 species with lower average abundance enriched in CRC patients. The divergence in species level pointed out the potential competition relationship among species. Another example is species *Akkermansia muciniphila* of phylum *Verrucomicrobia*. We noticed *Akk. muciniphila* is the only species that existed in samples of phylum *Verrucomicrobia*, meaning we should focus on the enrichment of *Akk. muciniphila* in healthy controls instead of its higher taxonomic levels. With the phylogenetic tree, taxonomy tree visualization sheds light on determining the responsible taxa of different abundance across taxonomic levels. (Fig.7A) As the taxonomy tree uses the average abundance of all samples to note the size of nodes, we further applied differential testing of all taxonomic levels. We used box plots to check the distribution of abundance along groups. We separately noted the significantly different taxonomy with *p* < p < 0.05 and p < 0.0001 with * and **. In the box plot of phylum and class, we detected that multiple outliers of both groups reflect the high heterogeneity within groups or caused by over a thousand samples involved (Fig.7B). The results can guide us in choosing a strict threshold to filter significantly different taxa. Finally, we could use the classifier module to build a gut microbiome-based CRC classifier with Random Forest. After the differential testing, we obtained 198 species with significant abundance differences between patients and healthy individuals. We chosen candidate biomarkers number by 10-fold cross-validation with five replicates and determine the optimal number of biomarker is six. (Fig.7C).

**Figure 7.**
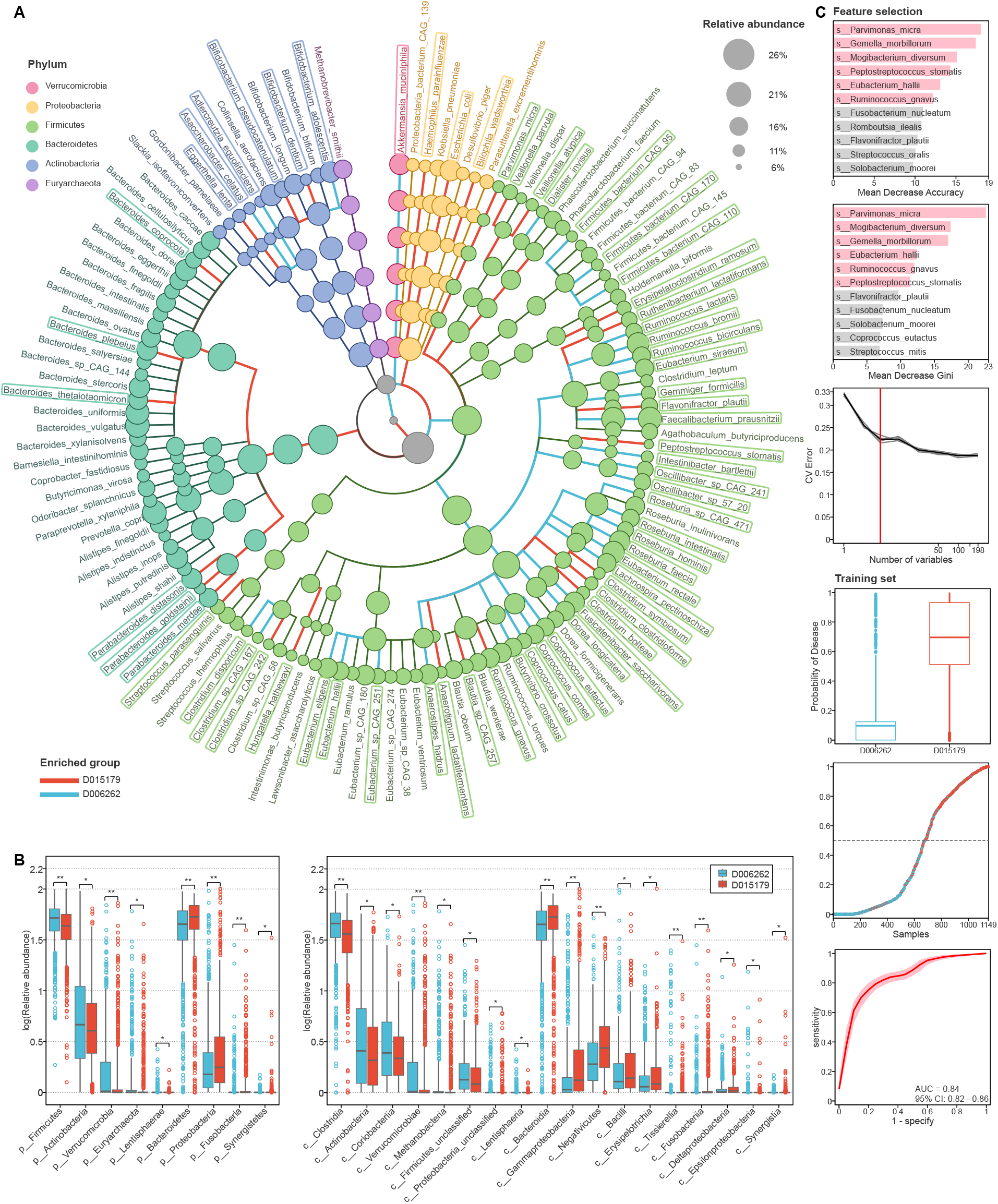
Microbiome composition comparison of samples from CRC patients and healthy individuals. **(A)** Taxonomy tree visualization of CRC dataset. **(B)** Differential testing visualizations in phylum and class. **(C)** Classifier visualization consists of feature selelction and evaluation in training set.

Based on the mean decrease accuracy and mean decrease Gini in the cross-validation process, we selected the top six species as candidate biomarkers to construct the random forest classifier. *Parvimonas micra*, *Gemella morbillorum*, *Mogibacterium diversum*, *Peptostreptococcus stomatis, Eubacterium hallii* and *Ruminococcus gnavus*. We further visualized the distribution of the probability of disease of training set samples in two groups in box plots and scatter plots. All in all, the classifier has AUC=0.84 with 95% CI from 0.82 to 0.86. The performance of the classifier is similar to the average performance of across-cohort predictions by Thomas *et al*. [6]. However, only *P. micra* and *P. stomatis* are the shared CRC biomarker species they also identified by independently univariate statistics.

The rest candidate biomarker species had been determined as CRC-related in the previous works except *M. diversum*. In early 2001, Ramon *et al.* reported the association of *G. morbillorum* and colon cancer [35]. A recent work of Yao *et al.* identified *G. morbillorum* as one of the non-invasive biomarkers of CRC from gutMEGA database [36]. *E. hallii* is a common gut microbe with probiotic properties regard to intestinal metabolic balance as contributing to propionate formation [37]. Several CRC-related gut microbiome research shows it has a significantly higher abundance in healthy individuals [38, 39]. *R. gnavus* is a well-known inflammatory polysaccharide producer associated with inflammatory bowel disease, and metabolic syndrome [40, 41]. The abundance of *R. gnavus* was reported to have a significant negative correlation with colorectal tumor numbers [42]. Mark *et al*. shown oral-source bacteria found in gut microbiome including, but not limited to, *P. stomatis*, *G. morbillorum*, and *P. micra*, related to the species diversity increasing and potentially relevant towards aiding tumor progression in CRC patients [43]. We identified *M. diversum* as a novel candidate CRC bacterial biomarker, an oral-source bacteria lacking functional experiments[44].

## Conclusions

We present a free access human gut microbiome WMGS sequencing data analysis platform GutMeta. To our knowledge, GutMeta is the first web server integrated database, online analysis, and visualization for the human gut microbiome. The built-in database of GutMeta, named HGDB, contains 20,989 WMGS sequencing samples with curated processed abundance and metadata. GutMeta provides implemented analysis modules containing alpha diversity, beta diversity, dimension reduction, enterotype, differential testing, and classifier. Every analysis modules have a corresponding interactive visualization developed on the GutMeta. We also designed two new visualizations to overview microbial composition among samples. Users can analyze their samples independently or co-analysis with a subset of HGDB on GutMeta by uploading their abundance and metadata. After the analysis task is finished, users can download their analysis results and visualize them online. GutMeta provides a one-stop web server for the researcher to analyze and visualize results and a reliable way to re-use public data.

## Availability and requirements

GutMeta is a publicly available Web platform accessible at https://gutmeta.deepomics.org/. Users are login-free to access and upload data to the platform.

## Availability of data and materials

Projects involved in the HGDB listed in https://gutmeta.deepomics.org/projects. Scirpts of analysis modules can be found in https://github.com/deepomicslab/GutMeta_analysis_scripts.

## Abbreviations

HGMD: Human Gut Metagenomics Database
WMGS: whole metagenome shotgun
SVG: Scalable Vector Graphics
IBS: irritable bowel syndrome
CRC: colorectal cancer
T2D: type 2 diabetes
MG: myasthenia gravis
IBD: inflammatory bowel disease
HMgDB: HumanMetagenomeDB
SRA: Sequence Read Archive
aov: analysis Of variance
UMAP: Uniform Manifold Approximation and Projection
PAM: Partitioning around medoids

## Declarations

Ethics approval and consent to participate Not applicable.

## Consent to publish

Not applicable.

## Author’s contributions

SCL conceived the study and coordinated the project. SCL and YJ designed the research. YJ collected the samples in the HGDB and implemented the analysis scripts. YW and LC developed the web server, which contains the front-end webpage, database, visualization, and back-end server. YJ and QZ prepared the abundance and metadata of samples in the HGDB. YJ and YF wrote the main manuscript text. SCL, YJ, YW, LC, and QZ reviewed the manuscript. All authors read and approved the final manuscript.

## Acknowledgements

We thank Hechen Li and Xikang Feng for assistance with Oviz and backend development.

